# The role of visual experience in brain inter-individual variability

**DOI:** 10.1101/2021.08.17.456515

**Authors:** Sriparna Sen, Ningcong Tong, Xiaoying Wang, Yanchao Bi, Ella Striem-Amit

## Abstract

Visual cortex organization is highly consistent across individuals. But to what degree does this consistency depend on life experience, in particular sensory experience? In this study, we asked whether visual cortex reorganization in congenital blindness results in connectivity patterns that are particularly variable across individuals, focusing on resting-state functional connectivity (RSFC) patterns from primary visual cortex. We show that the absence of shared visual experience results in more-variable RSFC patterns across blind individuals than sighted controls. Increased variability is specifically found in areas that show a group difference between the blind and sighted in their RSFC. These findings reveal a relationship between brain plasticity and individual variability in which reorganization manifests variably across individuals. We further investigated the different patterns of reorganization in the blind, showing that the connectivity to frontal regions, proposed to have a role in reorganization of the visual cortex of the blind towards higher cognitive roles, is highly variable. In a supplementary analysis, we link some of the variability in visual-to-frontal connectivity to another environmental factor – duration of formal education. Together, these findings show a role of sensory and socioeconomic experience in imposing consistency on brain organization. By revealing the idiosyncratic nature of neural reorganization, these findings highlight the importance of considering individual differences in fitting sensory aids and restoration approaches for vision loss.

**Significance statement:** The typical visual system is highly consistent across individuals. What are the origins of this consistency? Comparing the consistency of visual cortex connectivity between people born blind and sighted people, we showed that blindness results in higher variability, suggesting a key impact of individual experience on brain organization. Further, connectivity patterns that changed following blindness were particularly variable, resulting in diverse patterns of brain reorganization. Individual differences in reorganization were also directly affected by non-visual experiences in the blind (years of formal education). Together, these findings show a role of sensory and socioeconomic experiences in creating individual differences in brain organization and endorse the use of individual profiles for rehabilitation and restoration of vision loss.

## Introduction

The visual cortex has a highly-structured and consistent functional organization across individuals (Kanwisher, 2010; Wandell et al., 2007a). These include functional areas (e.g., those responding preferentially to faces, places, and body parts) in fixed cortical locations and consistent connectivity between them (Dehaene and Cohen, 2007; Gomez et al., 2018; Kanwisher, 2010; Kravitz et al., 2013; Weiner and Grill-Spector, 2013). Yet in addition to this broad invariance, some important variability in both activation patterns and connectivity is found across individuals (Feilong et al., 2018; Glezer and Riesenhuber, 2013; Osher et al., 2016; Saygin et al., 2011; Saygin et al., 2016; Tavor et al., 2016; Wang et al., 2015; Zhen et al., 2017; Zhen et al., 2015).

Brain variability informs theories of brain development and experience-dependent plasticity and also has clinical relevance. Sources of variability can be traced back to species-level developmental processes, showing that variability is greater in areas with greater evolutionary cortical expansion in humans, such as the parietal and frontal association cortices (Kaas, 2006; Mueller et al., 2013). Variability also hints at the temporal trajectory of development at the individual level (Gao et al., 2014), with a trajectory of changes accumulated differently across cortical sites and ages (Gao et al., 2014; Xu et al., 2018). Furthermore, variability between individuals in connectivity patterns and activation level has been linked to over a hundred behavioral abilities, from simple visual learning tasks (Baldassarre et al., 2012) to individuals’ traits and characteristics (e.g. (Tavor et al., 2016); reviewed in (Vaidya and Gordon, 2013)). In addition to the variability of the mature adult brain, individual differences have also been studied in development and aging, psychiatric illnesses, and developmental disorders (Brown, 2017; Foulkes and Blakemore, 2018; Friedman and Miyake, 2017; Hahamy et al., 2015). These individual differences are intensively studied to lead to better diagnosis and individually-tailored medical interventions (Drysdale et al., 2016; Fox and Greicius, 2010).

Despite the clear importance of inter-individual variability in determining brain development and (dys)function, the origins of neural variability remain unclear. Heritability has been shown to account for a high percentage of brain functional connectivity network organization (Ge et al., 2017; Reineberg et al., 2019; Yang et al., 2016) and brain functional activation (Alvarez et al., 2021; Park et al., 2012a; Park et al., 2012b; Polk et al., 2007), but does not explain the full range. One large source of variability, the effects of environmental factors such as sensory experience, remains particularly unclear. Unimodal cortices that develop fully early in life tend to show lower variability as compared to later-developing control and attention networks (Anderson et al., 2021; Mueller et al., 2013), suggesting a correlation between developmental trajectory and variability: A longer developmental trajectory allows for longer exposure to differential extrinsic experiences, causing higher variability in that brain region (Gratton et al., 2018; Mueller et al., 2013). But can experience also have a *stabilizing* effect on brain variability, in cases of shared environment and consistent experience? Is the low variability of the early cortices an inherent trait of these areas’ cortical tissue or is it due to the shared early-onset sensory experience in that modality? These questions broadly address the malleability of brain organization and the variability of potential outcomes when shared or typical experience is not provided.

Here we tested the role of experience on brain variability in an extreme model of experience deprivation: people born completely blind. In congenital blindness, the brain is deprived of the typical visual input that shapes the visual system (Maurer, 2017; Maurer et al., 2005; Röder et al., 2013; Wiesel and Hubel, 1963). We tested whether cross-individual variability in brain connectivity, manifested in resting-state functional connectivity (RSFC), is affected by sensory experience in a homogenous group of fully and congenitally blind adults. Although RSFC is only a correlate to functional responses and anatomical connectivity of the brain (Deco et al., 2011; Fox and Raichle, 2007; Honey et al., 2009; Smith et al., 2009), individual differences in connectivity appear to be stable across time (Badhwar et al., 2020; Jovicich et al., 2016; Liu et al., 2017), allowing their use for addressing questions of individual variability. Three possible predictions can be formulated with regard to the effect of sensory experience. On one hand, if visual experience serves to drive individual differences (as other life experiences have been proposed to do; (Gratton et al., 2018; Mueller et al., 2013)), limited visual experience in blind individuals may result in reduced inter-individual variability as compared to the sighted. On the other hand, the statistical properties of environmental experience in vision are highly consistent (Berkes et al., 2011; Simoncelli, 2003); this fact proposes an alternative hypothesis: consistent, structured visual input may have a stabilizing effect on brain variability. This stabilization would lead to higher RSFC diversity among blind individuals than among sighted individuals because their visual cortex organization is not constrained by shared visual experience. Mechanistically, this stabilization may stem from developmental pruning of variable non-visual projections innervating V1 (Dehay et al., 1984; Innocenti et al., 1988; Innocenti and Clarke, 1984; Kennedy et al., 1989; Rockland and Van Hoesen, 1994), and enforcing a more consistent connectivity profile. Either of these outcomes would highlight a significant role of experience in shaping individual neural differences. Yet a third alternative is that blindness would have no effect on brain consistency; this pattern would indicate strong inherited stabilization of brain individuation for the visual cortex that is not affected by experience.

Our analyses support the second hypothesis: Variability in connectivity from visual cortex is higher in blind individuals than in sighted individuals, suggesting a role of shared experience in promoting consistency of neural organization. Given this pattern, we further asked whether the plastic reorganization of visual cortex functional connectivity (Abboud and Cohen, 2019; Deen et al., 2015; Liu et al., 2007; Striem-Amit et al., 2015; Wang et al., 2013; Yu et al., 2008) manifests in a stereotypical, similar change across blind individuals, or if it is idiosyncratic. The blind visual cortex has been shown to respond, on average, to a large variety of sensory inputs and tasks across sensory modalities (Amedi et al., 2003; Bedny, 2017; Burton et al., 2003; Striem-Amit and Amedi, 2014; Striem-Amit et al., 2012a) and to become more functionally-connected to non-visual frontal cortices (Abboud and Cohen, 2019; Burton et al., 2014; Deen et al., 2015; Hawellek et al., 2013; Liu et al., 2007; Striem-Amit et al., 2015; Wang et al., 2013; Yu et al., 2008). However, the consistency of this reorganization has never been explicitly examined. Importantly, if reorganization varies among the blind, it could have far reaching implications towards implementing individually-tailored medical and rehabilitative interventions, as now explored for other disorders (Drysdale et al., 2016; Fox and Greicius, 2010), to address the large variability in sight restoration outcomes (Carlson et al., 1986; Ganesh et al., 2014; Gregory and Wallace, 1963; Huber et al., 2015).

## Results

We tested whether visual deprivation leads to altered inter-individual variability in the connectivity patterns of the visual cortex in a large group of congenitally fully-blind adults (n=25; see **Table 1** for the characteristics of the participants) and sighted adults (n=31) from two experimental cohorts scanned previously ((Striem-Amit et al., 2015; Striem-Amit et al., 2018); each cohort contained a blind and matched sighted group). We computed resting-state functional connectivity (RSFC) from an anatomically-defined seed in retinotopic primary visual cortex (V1), based on a visual localizer in an independent group of sighted individuals (Striem-Amit et al., 2015). As the cohort differences were negligible and highly localized (see **Fig. S1B,C**), RSFC maps across cohorts within each group (blind, sighted) were analyzed for their voxel-wise variability across individuals. To assess whether RSFC variability effects are indeed due to the absence of shared experience, the same procedures were computed for control seed regions in all non-visual Brodmann areas.

### V1 variability differs between congenitally blind and sighted individuals

We first tested whether there are differences in inter-individual variability of the V1-seeded RSFC resulting from blindness. For this aim, RSFC maps were analyzed using ANOVA (cohort × group; to remove any cohort effects, in addition to relevant preprocessing steps; see methods for detail; for main effects see **Fig. S1**). We calculated a whole-brain voxel-level homogeneity of variance test (Brown-Forsythe test (Brown and Forsythe, 1974); see detail in methods) for the group main effect, testing whether the two groups differed in their inter-individual variability of the RSFC values. This analysis revealed multiple areas that exhibit a significant inter-subject difference in V1-seeded RSFC variability between the blind and sighted groups (**Fig.1A**; group variability difference). These included areas of the ventral and dorsal visual pathways, posterior inferior parietal cortex and the inferior frontal cortex. Therefore, visual experience affects brain consistency. This analysis reveals only a non-directional difference in variability; to directly test the sign of the group difference, we calculated the ratio of variability between the groups (blind / sighted) across the brain (**Fig 1B**; ratio shown within areas that differ in variability between the groups). It is apparent that the blind show higher variability than the sighted in multiple areas, including parietal and frontal regions, with lower variability in only one small cluster in the right auditory cortex. Thus, visual experience can have an overall stabilizing effect on RSFC, and visual deprivation results in overall more variable RSFC from the visual cortex.

**Fig. 1.**
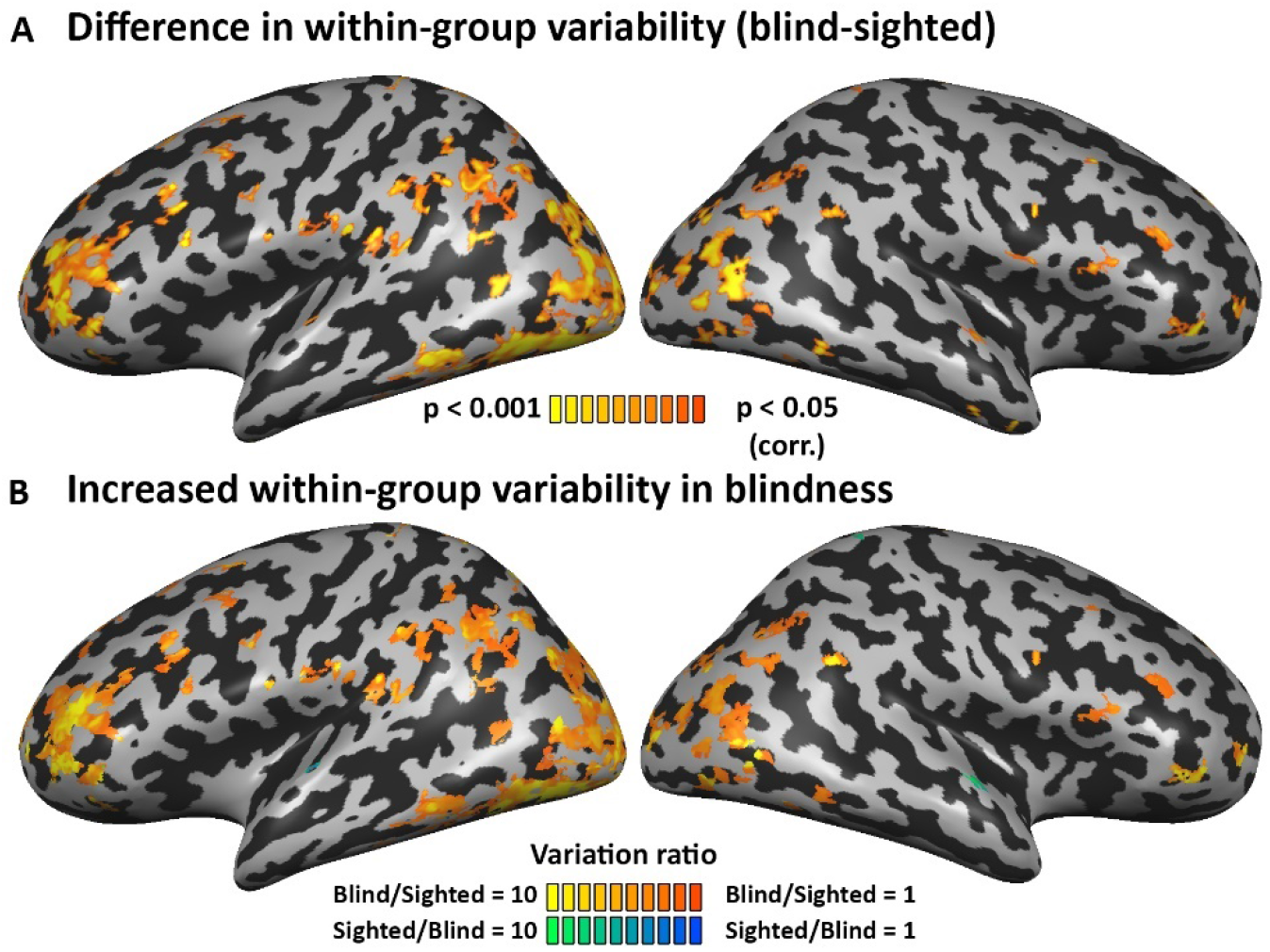
Variability in brain connectivity is increased in blindness. (**A**)The difference in within-group variability between the groups is significant in various parts of the brain, including in the frontal lobe. (**B**)Directional comparison of the within-group variability difference (ratio of blind intra-group variability divided by sighted intra-group variability > 1) shows that the blind have increased variability in most of the regions differing in their variation between the groups. This suggests a stabilizing effect of visual experience on visual cortex developmental functional connectivity.

### V1 variability increases especially for areas which reorganize in blindness

Inspecting inter-individual variability also allowed us to test if neural *reorganization* is consistent across blind individuals. Are areas whose connectivity and function have reorganized due to blindness also highly variable among blind individuals, as compared with the typical inter-individual differences for these areas? We tested this by inspecting the intra-group variability difference in the areas showing a main effect of group in the V1-RSFC values; areas showing change in V1-seeded RSFC between the blind and sighted (a 2-way ANOVA main group effect; **Fig. S1;** cohort and group × cohort interaction show very limited effects).

In accordance with previous work (Burton et al., 2014; Liu et al., 2007; Qin et al., 2014; Striem-Amit et al., 2015; Wang et al., 2013; Yu et al., 2008), blind individuals showed increased functional connectivity to some regions in the visual cortex, and several areas in the frontal lobe, including the inferior frontal sulcus (**Fig. 2A**). Given the proposal that increased connectivity with the frontal cortex drives reorganization in the visual cortex of the blind (Abboud and Cohen, 2019; Bedny, 2017; Deen et al., 2015; Rimmele et al., 2019), we focused on RSFC variability in these foci within the group of blind participants. Inferior frontal areas that show increased RSFC in the blind show double the variability within the blind group as within the sighted group (**Fig. 2B**; S^2^_sighted_ = 1.75, S^2^_Blind_ = 3.68).

**Fig. 2.**
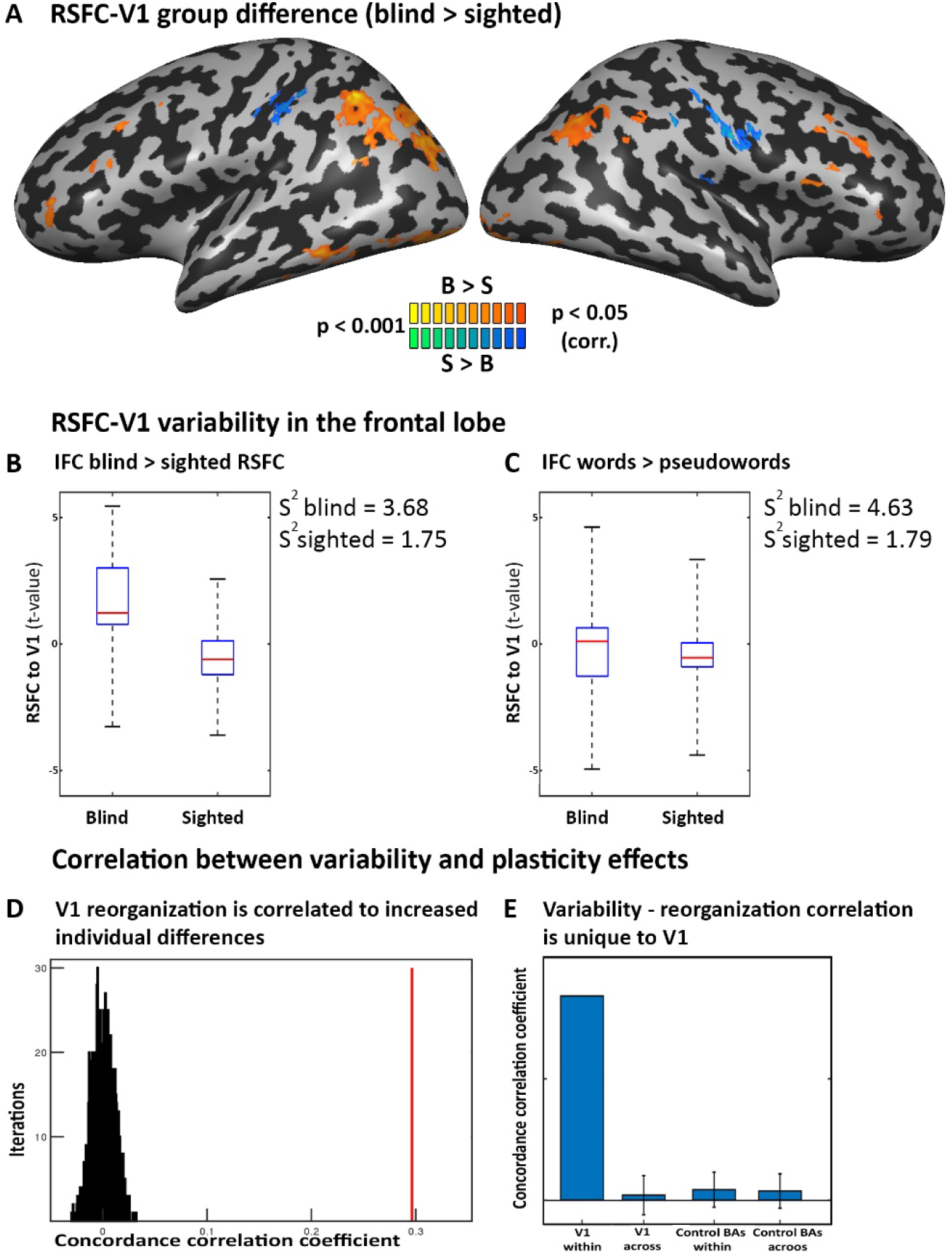
Brain reorganization in blindness is associated with increased inter-individual variability. (**A**) Increased V1-seeded RSFC in blindness is found in the visual streams, as well as in the bilateral inferior frontal cortex (IFC). (**B-C**) The blind show increased variability in their V1-seeded RSFC to left IFC frontal areas. **B**: within the areas showing increased RSFC in the blind (shown in Fig.2A); **C**: sampled in a language-selective IFC ROI, defined by preference towards words as compared to pseudowords. (**D**) Overall across the brain, areas showing changes in RSFC in blindness also show increased variability across blind participants. Concordance correlation coefficient was calculated between the RSFC group difference and RSFC change in variability for the V1 seed (red line) and compared to a spatial permutation test (distribution in black). (**E**) The link between reorganization and increased variability in blindness is specific to V1. Correlation between the two maps for the same seed was significantly greater than in correlating across seeds, and significantly greater for V1 as compared to other nonvisual Brodmann areas.

To specifically test frontal regions proposed to affect visual cortex reorganization, we also examined the variability of connectivity in left-lateralized frontal language regions. A spoken language-selective region was defined in the left inferior frontal sulcus (from a contrast of heard object names > heard pseudowords in a joint group of blind and sighted subjects from (Striem-Amit et al., 2018); see detail in methods). In this region as well, the intra-blind RSFC-with-V1 variability was more than double the intra-sighted variability (S^2^_sighted_ = 1.79, S^2^_blind_ = 4.63; see **Fig. 2C)**. Therefore, it appears that reorganization in the connectivity between the visual and frontal cortex in the blind is highly variable among the blind individuals.

Is this a general pattern, that neural reorganization manifests more variably in blindness? We correlated the spatial pattern of the group difference in mean RSFC from the visual cortex seed (**Fig. 2A**) with the variability difference between the groups (**Fig. 1A;** computed within areas showing RSFC with V1 in either group). The concordance correlation coefficient between the two maps (Lin, 1989) was highly significant (CCC=0.297, *p* < .0001; using a permutation test shuffling the voxels’ order, 100,000 iterations; **Fig 2D**). Therefore, it appears that when the brain reorganizes, it introduces a further source of variance, resulting in more diverse connectivity values. Importantly, the link between reorganization and variability is not an artefact due to the higher mean-difference between the groups: Using group-normalized RSFC values shows that the variability is increased in the blind even when controlling for the higher group mean value (**Fig. S2A,B)**. Similarly, the correlation between variability difference and reorganization remains when the variability difference is calculated from group-normalized data (CCC= 0.38, *p* < .0001; **Fig. S1C**).

Next, we tested the specificity of the link between reorganization and increased variability. If this pattern is driven by visual deprivation, we expected it to be especially prominent for the visual cortex seed, compared to seeds in non-visual areas. As a control, we performed the above analysis for seeds in each of the non-visual Brodmann areas, which did not show the same phenomena. Specifically, the correlation between variability and reorganization (between a Brown-Forsythe map and the ANOVA main effect of group for RSFC values) was significantly lower for other regions-of-interest (ROIs) than for V1 (V1 CCC within gray matter mask = 0.17, *p* < 0.001, non-visual Brodmann areas CCC = 0.009 ± 0.014 standard deviation, *p* < 0.208; for values for each BA separately see **Fig. S3**). Further, the cross-seed correlation, correlation between the group difference for V1 and the variability difference of any other Brodmann area; computed in a gray matter mask, was close to zero (CCC= 0.004; **Fig. 2E**), showing that the link between variability and reorganization is spatially specific. Therefore, it appears that the link between increase in variability and change in RSFC in the blind is linked specifically to connectivity with the visual cortex, suggesting that plasticity is characterized by increased variability and not by a ubiquitous change for all individuals.

### Spatial patterns variability across blind individuals

What forms does this increased variability take? To inspect if variability also manifests in different spatial patterns of connectivity in the blind, we used hierarchical clustering to group the blind individuals into clades based on their RSFC patterns and examined the RSFC pattern characterizing each subclade. This approach revealed informative diversity in the profiles of RSFC of the visual cortex among the blind individuals (**Fig. 3A;** see **Fig. 3B** for correlation matrix underlying this clustering). Most of the blind individuals clustered together in a clade showing (on average) focused positive RSFC with small foci in the inferior frontal cortex (clade 3; 17 individuals), along with differential patterns of RSFC with the superior frontal lobe (IFL; positive and negative values across individuals in different subclades (e.g. subclades III and IV). Curiously, in most of these subclades RSFC to the IFL was bilateral (subclades IV and V), whereas in a subclade of 7 individuals the pattern seemed lateralized to the left IFL (subclade VI in **Fig. 3A**). Two additional smaller clades seemed to cluster separately based on RSFC with the sensorimotor cortex – with a small clade (clade 2; 6 individuals) showing negative RSFC (anticorrelation) with the sensorimotor cortex, and two individuals (clade 1) showing a pattern of positive RSFC with the sensorimotor cortex as well as the inferior frontal cortex. Interestingly, the clustering did not show any qualitative distinction based on blindness etiology (**Fig. S4**), including a sparse distribution among clades for individuals whose blindness stemmed from genetic causes such as microphthalmia. Together, this analysis revealed a diverse pattern of organization relative to the visual cortex, across blind individuals.

**Fig. 3.**
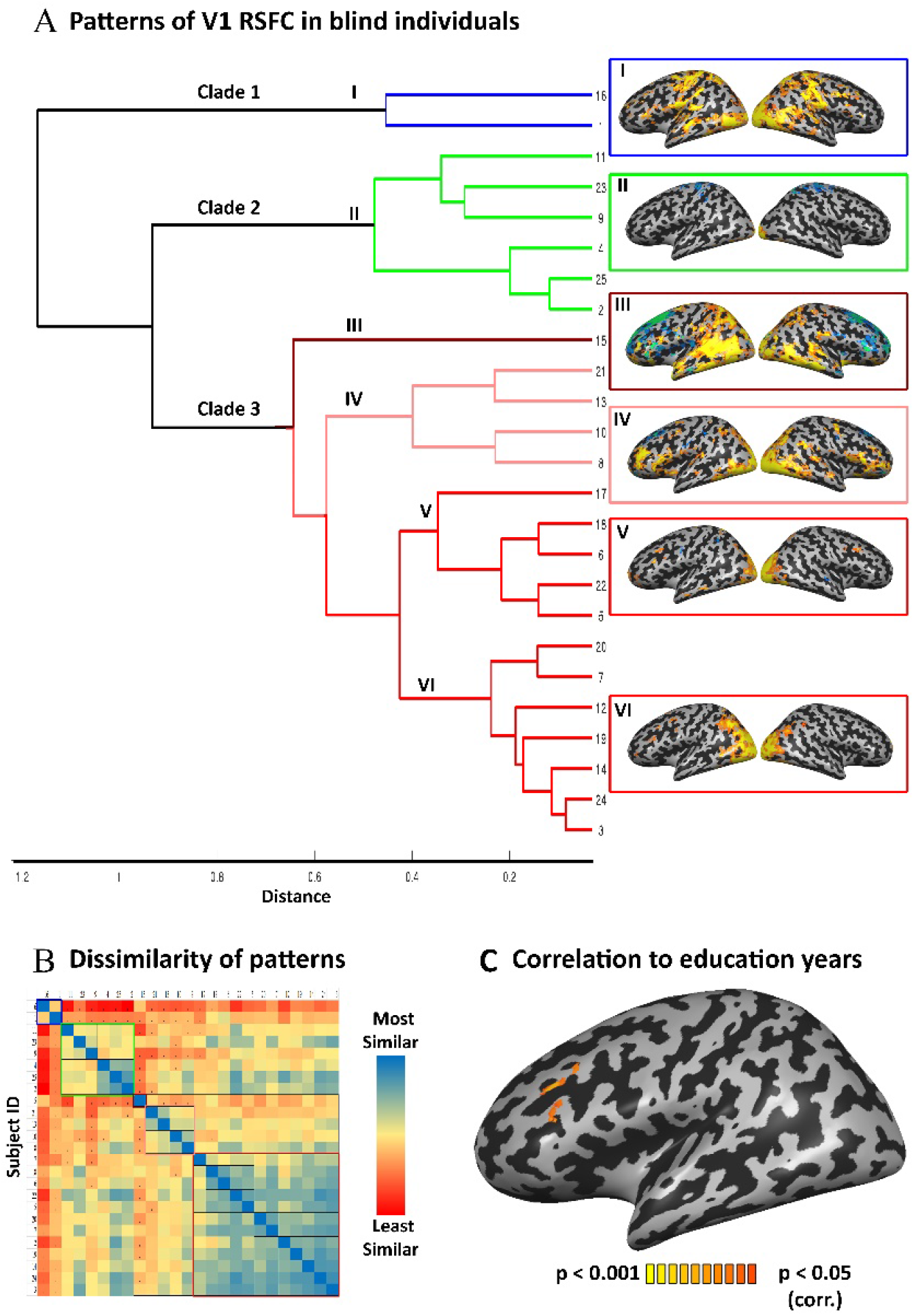
Patterns of brain reorganization in blindness. (**A**) V1-RSFC of each individual blind participant to each Brodmann area was used to compute hierarchical clustering of RSFC patterns across the blind. 3 main clades emerge, with differential connectivity to sensorimotor and frontal cortices. Subclades are marked with Roman numerals, and an average V1-RSFC map for the individuals in each subclade is shown. (**B**) The V1-RSFC correlation (dissimilarity) structure between individuals based on which the hierarchical clustering analysis was conducted. (**C**) V1-RSFC to the left-lateralized inferior frontal cortex in the blind (and not in the sighted) is correlated to the duration of formal education, showing one environmental factor affecting individual differences in brain reorganization in blindness.

Can we identify specific environmental factors contributing to this spatial diversity across blind individuals? As a supplementary analysis, we computed the correlation between V1-seeded RSFC and one socioeconomically-dependent factor: each individual’s years of formal education. We anticipated that visual cortex connectivity may be influenced by this factor because the visual cortex of the blind has been implicated in language (Abboud and Cohen, 2019; Amedi et al., 2004; Bedny et al., 2011; Burton et al., 2003), memory (Abboud and Cohen, 2019; Amedi et al., 2003; Raz et al., 2005), numerical thinking (Kanjlia et al., 2016) and executive function (Abboud and Cohen, 2019; Deen et al., 2015), all functions that are trained in formal education. Indeed, this was the case: V1-seeded RSFC with a region in the left inferior frontal cortex (dorsolateral prefrontal cortex) was correlated with education years in the blind group (**Fig. 3C**; peak Talairach coordinates −34, 28, 28). The sighted showed no correlation between years of education and RSFC from V1 to any brain region, including in the IFS clusters showing such correlation in the blind (sampled as a ROI in the sighted; r = 0.077, *p* = .68). Curiously, the IFG area which showed correlation to education duration is found in close proximity to areas showing increased variability between the blind and sighted, as well as increased RSFC in the blind group as compared to the sighted. Therefore, this exemplifies an interaction of blindness with environmental life circumstances that affects the diversity of visual cortex reorganization.

## Discussion

Inter-individual differences in brain organization stem from both hereditary and environmental factors. Here we examined the role of one extreme environmental factor, lack of visual experience, on the variability of the functional connectivity with the visual cortex. We showed that inter-individual differences in connectivity are higher in blind individuals (**Fig. 1B**), suggesting that shared sensory experience enforces consistency across individuals and brain network variability is expanded in its absence. Furthermore, we found that areas showing reorganization due to blindness, manifesting as increased RSFC with V1, also showed increased variability among blind individuals. This was true for specific areas in the frontal cortex language network proposed to underly reorganization in the blind and for the whole-brain pattern of reorganization (**Fig. 2**). This intra-group variability suggests that plasticity is not uniform among the blind, generating more variable outcomes than is typical in sighted individuals. Additionally, we qualitatively demonstrated some different spatial patterns that variable reorganization may take, by characterizing reorganization in distinct subgroups of blind individuals (**Fig. 3A**). While functional connectivity to the frontal lobe is a key characteristic of plasticity in blindness, we found that only some blind individuals show this pattern. Functional connectivity between the visual cortex and inferior frontal cortex (potentially related to working memory; (Rottschy et al., 2012)) is correlated with the duration of formal education, supporting a role for not only sensory but also social-educational factors in acquiring brain variability in blindness (**Fig. 3C**). These findings inform the developmental origins of individual variability, the properties of brain plasticity, and the importance of considering variability for rehabilitation of visual loss. In the next sections, we address all these topics in more depth.

The study of individual differences in brain reorganization has recently bloomed; inspecting identifiable “individual fingerprints” in brain connectivity and their behavioral correlates (Finn et al., 2015; Gratton et al., 2018), has been aided in particular by large data-collection initiatives (Biswal et al., 2010; Di Martino et al., 2014; Smith et al., 2013). These differences have been found not only in correlations with specific behavioral skills (Baldassarre et al., 2012; Finn et al., 2015; Fong et al., 2019; Koyama et al., 2011; Vaidya and Gordon, 2013; Wang et al., 2016), but also in broader applications. Individual differences have been proposed as a basis for quantitative phenotypes and biomarkers to be integrated into molecular and genetic studies of human neurological and psychiatric diseases (Biswal et al., 2010; Rosenberg et al., 2015; Xin et al., 2019) and to guide medical interventions (Drysdale et al., 2016; Fox and Greicius, 2010). Importantly, individual differences in connectivity appear to be stable across time (Badhwar et al., 2020; Jovicich et al., 2016; Liu et al., 2017), suggesting they reflect true anatomical and functional differences, rather than merely different temporary cognitive states during the scans. However, the contributing factors underlying this variability are not clear. A role for inherited genetic components of neural variability is evident (Anderson et al., 2021; Gao et al., 2014; Ge et al., 2017; Jansen et al., 2015; Koten et al., 2009; Park et al., 2012a; Park et al., 2012b; Polk et al., 2007; Thompson et al., 2001; Xin et al., 2019); a recent paper has even found specific genetic components implicated in brain plasticity underlying variability in functional connectivity of multisensory integration areas in blind children (Ortiz-Terán et al., 2017). However, there are complex nonlinear interactions between inherited components and age/maturation across different cortical systems (Gao et al., 2014), requiring a better understanding of the environmental components. Developmental studies have examined the role of social-environmental factors such as socioeconomic status (Foulkes and Blakemore, 2018) in inter-individual differences, highlighting the adverse effects of socioemotional deprived environments on development in children and adolescents (e.g. (Gunnar and Reid, 2019; Herzberg and Gunnar, 2020)). The results of these studies emphasize the potential benefits of understanding plasticity through the lens of individual differences.

Here we studied the role of a more extreme form of environmental change: complete deprivation of an entire sensory channel. Using this model, aided by an unparalleled cohort-size for this homogenous unique population, we showed that experience has immense effects on individual differences and can modify the variability in the neural connectivity profile of extensive cortical tissue. In the past, functional connectivity variability was found to be highest in association cortices that developed phylogenetically recently (Kaas, 2006; Krubitzer and Prescott, 2018; Smaers et al., 2011), such as the parietal, temporal and frontal association cortices, whereas early sensory cortices exhibited low individual differences (Anderson et al., 2021; Fischl et al., 2007; Mueller et al., 2013; Xu et al., 2018). However, this divide also exists within the same areas’ ontogenetic developmental trajectory; these high-cognitive function and association networks continue to develop through adolescence, whereas lower sensorimotor systems reach maturity earlier (Amlien et al., 2014; Guillery, 2005; Raznahan et al., 2011; Shaw et al., 2008; Xu et al., 2018). Therefore, the study of typically-developed individuals does not allow us to resolve whether individual variability in these regions results from longer exposure to environmental factors in the individual’s lifetime or evolutionary diversity. Our study shows how experience can affect even a relatively evolutionarily-conserved and typically highly-consistent cortical area, whose connectivity typically stabilizes in early childhood (Xu et al., 2018). Furthermore, we showed how a social environmental factor, years of education, which extends into adulthood, affects variability of RSFC in V1. Thus, even in the case of the early visual cortex, experience over long timescales can enhance individual differences, disentangling the roles of phylogenetic and ontogenetic development on brain organization.

Our findings suggest a link between variability and plasticity in brain development. Not only was the visual RSFC more variable in the blind, but the variability was specifically increased in areas that showed reorganization due to blindness. Though this is a correlational finding, it seems plausible that the absence of otherwise-consistent experience would remove potential constraints on development, allowing more variability between individuals. This change might take place especially during brain development stages in which fine-tuning of cortical structure and anatomical connectivity is done. In other mammals, these include stages of pruning of exuberant connectivity, which is based in part on activity-dependent patterns (Innocenti and Price, 2005). Therefore, as suggested previously (Amedi et al., 2003; Collignon et al., 2009; Sathian, 2005), transient connectivity to the visual cortex (Dehay et al., 1984; Innocenti et al., 1988; Innocenti and Clarke, 1984; Kennedy et al., 1989; Rockland and Van Hoesen, 1994) that is typically pruned following visual experience may endure in blind humans, to variable extents across individuals (thus not necessarily apparent in group-level analyses; (Fine and Park, 2018)). Pruning change as a result of visual or sensory experience is seen in other mammalian species (Henschke et al., 2017; Karlen et al., 2006; Nicolelis et al., 1991), and in non-human primates the absence of visual experience can also cause changes to corticogenesis (Magrou et al., 2017). An alternative but non-exclusive account is that the variability reflected in the RSFC networks shown here stems from shorter-term changes in brain connectivity, such as those associated with unmasking of existing but otherwise dormant connections between the visual and non-visual cortices (Hamilton and Pascual-Leone, 1998; Rauschecker, 1995). Although in the absence of a late-onset blindness group it is impossible to fully discern these two accounts, many studies have demonstrated that plasticity in early visual cortex in late-onset blindness is greatly reduced as compared to congenital blindness (Burton et al., 2002a; Burton et al., 2002b; Carlo et al., 2019; Cohen et al., 1999; Collignon et al., 2013; Fujii et al., 2009; Wittenberg et al., 2004), suggesting that processes beyond unmasking are involved in generating non-visual responses and RSFC in the congenitally blind. Regardless of the underlying mechanism, this data shows that plasticity allows an increase in the breadth of potential outcomes for brain organization.

What are the sources of the differential variability between the blind and the sighted? In terms of visual experience, the blind participants are a homogenous group of congenitally and fully-blind adults, without any ability to recognize visual shapes. The change in variability in brain connectivity we found can therefore not be linked to different levels of visual experience. Although we cannot exclude that the origins of some of these differences may be genetically linked to the causes of blindness, it is worth noting that only some of the participants’ blindness stemmed from clearly heritable conditions such as microphthalmia (Bardakjian and Schneider, 2011), and even in these cases the spatial profiles of connectivity did not seem to cluster based on blindness etiology (including for siblings; see **Fig. S4**). Instead, variability may be ascribed to two sources. The first is the absence of the typical visual input, which is characterized by specific and similar statistical properties (Berkes et al., 2011; Simoncelli, 2003). It is well known that visual experience influences brain organization and function (Arcaro et al., 2017; Cloherty et al., 2016; Dehaene et al., 2010; Espinosa and Stryker, 2012; Golarai et al., 2017; Gomez et al., 2019; Hubel and Wiesel, 1964; Maurer et al., 2005; Ostrovsky et al., 2006; Röder et al., 2013; Ruthazer and Aizenman, 2010; Sugita, 2004; Sugita, 2008; Wiesel and Hubel, 1963). As the visual system properties are evolutionarily tailored to the environment statistical properties (Simoncelli and Olshausen, 2001), confirmatory and typical external experience may strongly enforce typical organization and connectivity strengths: decreasing and pruning the less-dominant, and otherwise partly transient non-visual inputs (Dehay et al., 1984; Dehay et al., 1988; Innocenti and Clarke, 1984; Innocenti and Price, 2005; Kennedy et al., 1989; Rockland and Ojima, 2003; Rockland and Van Hoesen, 1994) (e.g., from frontal cortices) which may be more variable across individuals, and resulting in a relatively consistent system. A lack of a shared experience may lead to increased inter-individual variability in the blind, as (likely already variable) non-dominant inputs, variable in themselves, may be strengthened by small environmental experiences, genetic predispositions or random noise, driving different individuals to different strengthening of FC with different systems.

Another source of variability which is not mutually exclusive, could be individual adaptations to blindness, such as the compensatory use of other senses (Beaulieu-Lefebvre et al., 2011; Collignon et al., 2009; Goldreich and Kanics, 2003; Röder et al., 1999; Van Boven et al., 2000) and cognitive faculties (e.g., increased reliance and improved memory and verbal skills (Dormal et al., 2017; Loiotile et al., 2019; Occelli et al., 2016; Pozar, 1982; Raz et al., 2007; Tillman and Bashaw, 1968)). Plasticity correlated to these different abilities has been found in the visual cortex of the blind (e.g. (Amedi et al., 2003; Gougoux et al., 2005)), and differential abilities and reliance on these modes of compensation (e.g., reading braille books as opposed to listening to audiobooks) across individuals could lead to variability in visual system connectivity. In our characterization of the increased variability of the blind, we are unable to separate these two accounts completely. In a partial attempt to do so, we have shown here that the RSFC of the visual cortex to the left inferior frontal cortex is correlated to an individual’s duration of formal education. However, most of the regions that showed changes in variability were not accounted for in this preliminary exploration. Furthermore, overall increased variability was not found in non-visual sensory areas (auditory and somatosensory cortices), making it unlikely that experience or expertise in compensatory senses underlies the full variability. Future work should parse out the effects of specific environmental and personal factors affecting the reorganization in the blind, beyond the lack of shared visual experience.

Based on our exploratory clustering analysis, reorganization generates not only variability in connectivity, but also distinct spatial connectivity profiles. For example, some blind individuals show strong positive and others negative connectivity between the visual and sensorimotor cortices. Most of the blind show RSFC between V1 and the inferior frontal cortex, but connectivity to the superior frontal cortex differs between subclades. Although a full characterization of individual profiles would benefit from additional correlates and an increased sample size, we can already gain two interesting insights. The first is that the most drastic form of reorganization associated with blindness, functional connectivity to the left frontal cortex (Burton et al., 2014; Liu et al., 2007; Qin et al., 2014; Striem-Amit et al., 2015; Wang et al., 2013; Yu et al., 2008), which has been described as related to language (Bedny, 2017), and thus may be expected to be lateralized to the left, is found to be lateralized only in a minority of the subjects (7 of 25 participants; subclade VI; **Fig. 3A**). Overall, the RSFC between V1 and frontal cortex is quite variable and often bilateral (**Fig. 2B,C**). An additional finding is that differences are found between individuals in connectivity to the somatosensory cortex: clades 1 and 2 showed positive and negative RSFC, respectively, to the sensorimotor cortex (**Fig. 3A**). This pattern suggests potentially-informative changes in the link between the senses and the importance of reorganization regarding touch in different blind individuals.

These different reorganization profiles may have clinical implications for vision rehabilitation. The causes of the high variability of outcomes of sight restoration attempts using cataract removal and stem-cell therapy (Carlson et al., 1986; Ganesh et al., 2014; Gregory and Wallace, 1963; Huber et al., 2015) are currently unknown, with some patients gaining little functional sight. As evident from cochlear implantation in deafness ((Feng et al., 2018; Lee et al., 2001; Olds et al., 2016), c.f. (Heimler et al., 2014; Land et al., 2016; Lyness et al., 2013)), variability in restoration for a missing sense may depend on whether the neural system for that sense is intact, or whether it has undergone cross-modal reorganization and is no longer capable of processing information of the original modality. Similar considerations may apply to visual restoration as well: Some of the failed sight restoration attempts may have neural causes (Striem-Amit et al., 2011). In contrast to these invasive methods that require an intact (rather than cross-modally dominant) visual system, assistive and adaptive technologies such as sensory substitution devices are designed to utilize cross-modal translations. For example, sensory substitution devices that convert visual images into sounds (Capelle et al., 1998; Meijer, 1992; Striem-Amit et al., 2012b) or touch (Bach-Y-Rita et al., 1969) benefit from individually increased levels of non-visual capacities (Arnold et al., 2017; Brown et al., 2011), thus potentially benefitting from cross-modal plasticity of specific senses. In late-onset vision loss due to age-related diseases (e.g., macular degeneration, glaucoma, cataracts) there is a dizzying selection of sensory aids and substitution techniques. For the task of reading alone, approaches include refreshable Braille displays, screen readers, optical (magnifying glasses) and electronic aids (video magnifiers), employing touch, audition, and vision respectively. Similar diversity exists for navigation needs (guide dog, white cane, electronic canes, smart glasses). It may be beneficial for individuals facing this array of options, whether they are adults facing vision loss or children born with visual disabilities, to have a suggestion of which of these technologies will be most effective based on their own neural plasticity profile. Understanding variability and individual differences in cross-modal reorganization levels (from anatomical markers, such as V1 thickness; (Aguirre et al., 2016) or functional connectivity differences as here) may allow for individually-tailored, personalized medicine and assistive technology in sight rehabilitation of visual disorders.

In conclusion, we showed that in the absence of sensory experience (due to blindness), brain reorganization generates larger inter-individual variability beyond the individual differences found in the typical sighted population. Variability is increased especially for areas that have reorganized in their connectivity to V1 due to blindness, and blind individuals show different spatial patterns of connectivity of their visual cortex. This finding suggests an important role for experience in determining the individual variability of neural organization. Additionally, these results highlight the need to consider idiosyncratic profiles of plasticity in tailoring rehabilitation plans for individuals with sensory deficits.

## Materials and methods

### Participants

Twenty-five congenitally-blind individuals and 31 sighted controls participated in the study. The data was collected for two previous studies (Striem-Amit et al., 2015; Striem-Amit et al., 2018), scanned at two separate sites. Cohort A included 13 congenitally-blind individuals and 18 sighted controls (Striem-Amit et al., 2015). Cohort B included 12 congenitally-blind individuals and 13 sighted controls (Striem-Amit et al., 2018). Sighted participants had normal or corrected-to-normal vision; all participants had no history of neurological disorder. Groups within each cohort were matched for age and education. Participants in the blind group (across cohorts) were between 22 and 63 years of age (no significant group difference for each cohort separately, *p* > .14, *p* > .99, or collapsed across cohorts *p* > .34). Duration of formal education was also comparable across groups (*p* > .45, *p* > .97 for each cohort separately, or collapsed across cohorts *p* > .49). See **Table S1** for detailed characteristics of the blind participants in each cohort. The Tel-Aviv Sourasky Medical Center Ethics Committee approved the experimental procedure for cohort A and the institutional review board of the Department of Psychology, Peking University, China and the institutional review board of Harvard University approved the experimental procedure for cohort B. Written informed consent was obtained from each participant.

### Functional Imaging

Functional magnetic resonance imaging (fMRI) data were obtained during resting conditions, without any external stimulation or task (i.e., spontaneous blood oxygen level-dependent fluctuations) for both cohorts. During the scan, subjects lay supine in the scanner with no external stimulation or explicit task. The sighted subjects were blindfolded and had their eyes shut for the duration of the scan.

Cohort A: Images were acquired with a 3-T General Electric scanner with an InVivo 8-channel head coil. Data was comprised of one functional run, containing 180 continuous whole-brain functional volumes acquired with an Echo Planar Imaging sequence (Repetition Time = 3000 ms, Echo Time = 30 ms, 29-46 slices, voxel size 3 × 3 × 4 mm, flip angle 90°, 182 volumes, scan length = 9.1 min). T1-weighted anatomical images were acquired using a 3D MPRAGE sequence (typical scan parameters were: 58 slices; TR = 8.9 ms; TE = 3.5 ms; inversion time = 450ms; FA = 13°; FOV = 256 × 256 mm; voxel size = 1 × 1 × 1 mm; matrix size = 256 × 256).

Cohort B: Images were acquired using a Siemens Prisma 3-T scanner with a 20-channel phase-array head coil. Data was comprised of one functional run, containing 240 continuous whole-brain functional volumes that were acquired with a simultaneous multi-slice (SMS) sequence supplied by Siemens: slice planes scanned along the rectal gyrus, 64 slices, phase encoding direction from posterior to anterior; 2 mm thickness; 0.2 mm gap; multi-band factor=2; TR = 2000 ms; TE = 30 ms; FA = 90°; matrix size = 112 ×112; FOV = 224 × 224 mm; voxel size = 2× 2×2 mm. T1-weighted anatomical images were acquired using a 3D MPRAGE sequence (192 slices; 1mm thickness; TR = 2530ms; TE = 2.98ms; inversion time = 1100ms; FA = 7°; FOV = 256 × 224 mm; voxel size = 0.5 × 0.5 × 1 mm, interpolated; matrix size = 512 × 448). Data of cohort B were down-sampled to a resolution of 3 mm iso-voxels for joint analysis with data from cohort A.

### fMRI preprocessing

Data analysis was performed using the BrainVoyager 20 software package (Brain Innovation, Maastricht, Netherlands) and custom scripts in MATLAB (MathWorks, Natick, MA) following standard preprocessing procedures. The first two images of each scan were excluded due to non-steady state magnetization. Preprocessing of functional scans included 3-Dimensional motion correction, slice scan time correction, band pass filtering (0.01-0.1Hz), regression of spurious signals from the ventricles and white matter regions (defined using the grow-region function in Brain Voyager on the individual level), and spatial smoothing with a 4-mm full-width-at-half-maximum (FWHM) Gaussian kernel. Head motion did not exceed 2 mm along any given axis or include spike-like motion of more than 1 mm in any direction. Data were normalized to standard Talairach space (Talairach and Tournoux, 1988). In order to overcome differences originating from the two datasets’ differences in scan parameters and cohorts, we applied post-hoc standardization (z-normalization of the data), shown to dramatically reduce site-related effects (Yan et al., 2013). An additional step to exclude site-related effects was the integration of the cohort grouping factor explicitly in the RSFC ANOVA (see below), and study effects related to group regardless of the cohort (as evident by the minimal cohort effects remaining in the analyzed data; **Fig.S1**).

### Seed regions-of-interest

The region-of-interest (ROI) for the primary visual cortex (V1) was defined from an independent localizer, acquired in a separate group of 14 sighted subjects (Striem-Amit et al., 2015) using a standard phase-encoded retinotopic mapping protocol, with eccentricity and polar mapping of ring and wedge stimuli, respectively (Engel et al., 1994; Sereno et al., 1995; Wandell et al., 2007b; Wandell and Winawer, 2011). The experimental detail can be found in (Striem-Amit et al., 2015). Polar mapping data were used to define the borders of V1, used as a seed ROI for the RSFC analyses. Non-V1 seed regions employed in the control analysis included anatomically-defined Brodmann areas (from the anatomical atlas in Brainvoyager) with the exception of visual areas BA 17, 18 and 19.

### RSFC variability analyses

Individual time courses from the V1 seed ROI were sampled from each of the participants and used as individual predictors in RSFC seed analyses. Data were analyzed with a 2 × 2 random effects ANOVA (Group [blind, sighted] × Cohort [A,B]) at the voxel level. In addition to the main effect of Group (**Fig.S 1A**, see limited cohort effect and group X cohort interaction in **Fig. S1B,C**), we calculated the Brown-Forsythe test for equal variance for this main effect, testing whether the two groups differed in their inter-individual variability of the RSFC values (**Fig. 1A**). The Brown-Forsythe test (Brown and Forsythe, 1974) is a homogeneity of variance test similar to the Levene test, conventionally used to test for variability differences, but utilizes the median instead of the mean, safeguarding against false positives in cases of skewed data distribution (Olejnik and Algina, 1987). The same analyses were performed for all non-visual control seed ROIs (Brodmann areas) for the comparison of variability and reorganization correlation (see detail below). The minimum significance level of all results presented in this study was set to *p* < .05, corrected for multiple comparisons within the gray matter volume using the spatial extent method (Forman et al., 1995; Friston et al., 1993) (a set-level statistical inference correction). Correction was based on the Monte Carlo simulation approach, extended to 3D datasets using the threshold size plug-in for BrainVoyager QX. We additionally computed the variability of RSFC within each group separately, using each group’s normalized data to overcome possible effects of the different cohorts on the mean and standard deviation of the RSFC. To inspect the direction of the variability group effect, we computed the ratio of variability between the groups (Variability_Blind_ / Variability_Sighted_; **Fig. 1B, Fig. S2B**) for each voxel showing a significant Brown-Forsythe test effect (*p* < .05, corrected). The same computation of the variability ratio was also conducted within left frontal ROIs (see detail below).

To inspect the direction of reorganization in V1 RSFC, in addition to the ANOVA model of the main effect of group on V1-RSFC (Fig. S1), we computed a post-hoc t-test comparing RSFC between the groups (blind vs. sighted).

To quantitatively assess the link between reorganization in the blind and variability effects, we compared the spatial pattern of variability (**Fig. 1A**) and reorganization in the blind (**Fig. S1**), by computing the concordance correlation coefficients (CCC; (Lin, 1989)) between these maps, within the gray matter. CCCs were computed using custom software written in MATLAB (MathWorks, Natick, MA). Concordance correlation coefficient values range from 1 (perfect spatial similarity) to – 1 (perfect spatial dissimilarity). While CCC, similarly to Pearson’s linear correlation coefficient, tests for shared fluctuations in variance of two datasets, it also penalizes for differences in means between the two sets, thus serving as a more sensitive measure for map differences in both spatial patterns and overall values. The significance level for the CCCs was obtained using a permutation test (100,000 iterations) randomly shuffling voxels from one map and convolving the resulting map with a Gaussian kernel based on data smoothness estimation, to account for spatial autocorrelation. As an additional control, we compared the CCC values across regions of interest, for pairs of maps (a variability map and a group-difference map) stemming from a coupled comparison for the same seed ROI, as compared to correlation values stemming from comparisons of variability and group-difference maps across seed ROIs. For example, computing the CCC between the Brown-Forsythe test map for the V1 seed and the map of blind-sighted group effect for the same seed, as compared to the CCC between the Brown-Forsythe test for the V1 seed and the map of blind-sighted group effect for each of the other non-visual Brodmann area seed ROIs.

### Variability ROI analysis

To inspect the variability of the frontal lobe, two sampling regions were used. (1) clusters in the inferior frontal lobe showing increased functional connectivity from V1 in the blind in the present study (seen in **Fig. 2A**) (2) a left lateralized language-selective region in the inferior frontal cortex (Talairach coordinates −29, 15, 18), defined from a contrast of heard object names greater than heard pseudowords in the joint group of blind and sighted subjects from cohort B. The full experimental protocol for this contrast is detailed in (Striem-Amit et al., 2018); briefly, auditory pseudowords and words from different concept categories were presented in a block-design fMRI experiment. For both ROIs, we sampled V1 RSFC GLM parameter estimates from each of the participants, and the variability (S^2^) of each group was calculated as well as the ratio between them, to assess whether the groups’ differed in their intra-group variabilities.

### Clustering analysis

To qualitatively explore individual differences in the RSFC from the visual cortex of the blind, we performed a hierarchical clustering analysis across subjects V1-seeded RSFC maps, using RSFC values for each individual from each of the Brodmann areas in the BrainVoyager atlas (see above). Distance was calculated as the correlation between individual RSFC vectors, implemented in MATLAB (MathWorks, Natick, MA). A dendrogram of the distances across all participants was computed based on complete distance between clusters (**Fig. 3A**; for the underlying correlation dissimilarity matrix see **Fig. 3B**). As a preliminary quantitative exploration of the clustering analysis, the average RSFC pattern (average V1-RSFC t map across the subjects) for individuals within each subclade was computed.

### Correlation with education

As a preliminarily analysis to inspect the effect of specific environmental factors on V1 RSFC variability, we calculated the correlation between each voxel’s V1-seeded RSFC for all participants with the number of years of formal schooling they received, for each group separately (**Fig. 2E** for the blind; the sighted showed no significant correlation at *p*<0.05 corrected). In the IFS cluster showing such correlation in the blind, correlation in the sighted group was also sampled, and was found to be non-significant (*p* = .68).

## Acknowledgements

We are thankful to the blind subjects who participated in our experiment, to Amir Amedi for invaluable contribution to the work, to Josef P. Rauschecker and Maeve Barrett for helpful discussions and comments and to Hila Zadka and Ao Yuan for statistical advising. This work was supported by the National Natural Science Foundation of China (31500882 to X.Y.W., 31671128 to Y.B.); the Fundamental Research Funds for the Central Universities (2017XTCX04, to Y.B.) and Interdisciplinary Research Funds of Beijing Normal University (to Y.B.) and the Edwin H. Richard and Elisabeth Richard von Matsch Distinguished Professorship in Neurological Diseases (to ESA).

## Supplementary Material

**Table S1:** Characteristics of blind participants.

| Subject | Cohort | Gender | Age | Cause of blindness | Light perception | Handedness | Age of blindness onset |
| --- | --- | --- | --- | --- | --- | --- | --- |
| 1 | A | F | 29 | Microphthalmia | None | Right | 0 |
| 2 | A | F | 23 | Microphthalmia, retinal detachment | None | Left | 0 |
| 3 | A | F | 30 | Retinopathy of prematurity | None | Right | 0 |
| 4 | A | M | 37 | Retinopathy of prematurity | None | Right | 0 |
| 5 | A | F | 38 | Enophthalmus | None | Left | 0 |
| 6 | A | M | 54 | Retinopathy of prematurity | None | Right | 0 |
| 7 | A | M | 23 | Microphthalmia | None | Right | 0 |
| 8 | A | F | 34 | Retinopathy of prematurity | None | Right | 0 |
| 9 | A | M | 31 | Retinopathy of prematurity | None | Right | 0 |
| 10 | A | F | 35 | Retinoblastoma | None | Right | 0 |
| 11 | A | F | 34 | Microphthalmia | None | Left | 0 |
| 12 | A | F | 30 | Leber congenital amaurosis | Faint | Ambidextrous | 0 |
| 13 | A | M | 42 | Retinopathy of prematurity | Faint | Right | 0 |
| 14 | B | M | 36 | Microphthalmia | None | Ambidextrous | 0 |
| 15 | B | M | 22 | Microphthalmia | None | Right | 0 |
| 16 | B | M | 33 | Microphthalmia; microcornea | None | Right | 0 |
| 17 | B | M | 48 | Glaucoma | None | Right | 0 |
| 18 | B | F | 46 | Glaucoma | None | Right | 0 |
| 19 | B | M | 40 | Leukoma | Faint | Right | 0 |
| 20 | B | F | 50 | Cataracts; eyeball dysplasia | Faint | Right | 0 |
| 21 | B | M | 57 | Eyeball dysplasia | None | Right | 0 |
| 22 | B | F | 43 | Glaucoma | None | Right | 0 |
| 23 | B | M | 48 | Microphthalmia; cataracts; leukoma | None | Right | 0 |
| 24 | B | M | 63 | Glaucoma; leukoma | None | Right | 0 |
| 25 | B | F | 41 | Optic nerve atrophy | Faint | Right | 0 |

\* Cohort A was acquired in Israel and comprised of 13 blind adults and 18 sighted controls (Striem-Amit et al., 2015).
Cohort B was acquired in China and comprised of 12 blind adults and 13 sighted controls (Striem-Amit et al., 2018).

**Figure S1:**
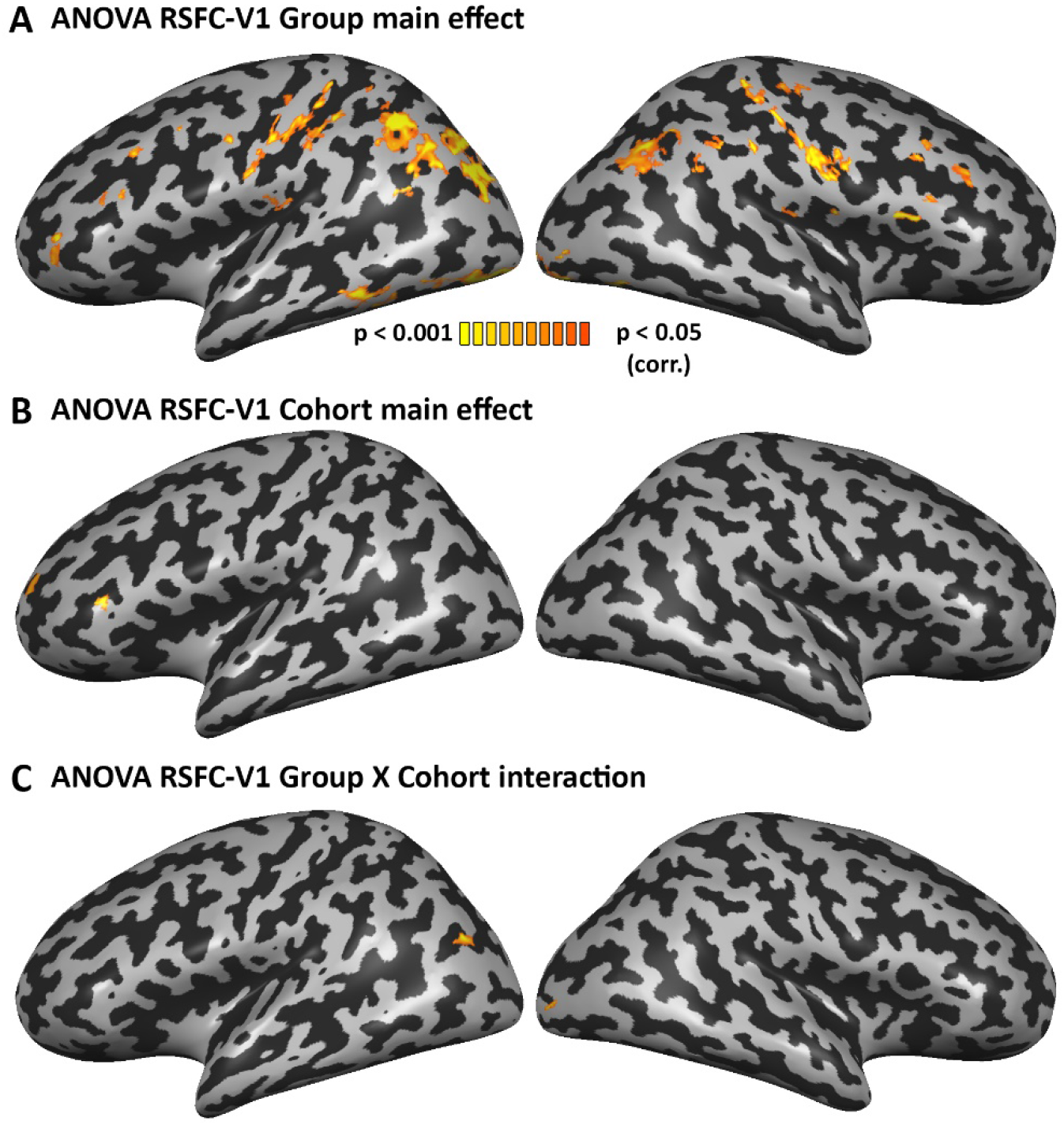
Main effect of sight. A. A main effect of sight across the cohorts is depicted. As reported before, the blind and sighted differed in their RSFC from the visual cortex to visual, parietal and frontal regions. A direct contrast between the groups is depicted in **Fig. 2A**. B. The main effect of cohort across the groups, showing little difference focused in the left inferior frontal (orbitalis) cortex. C. The Group X Cohort interaction shows little early visual cortex effect.

**Figure S2.**
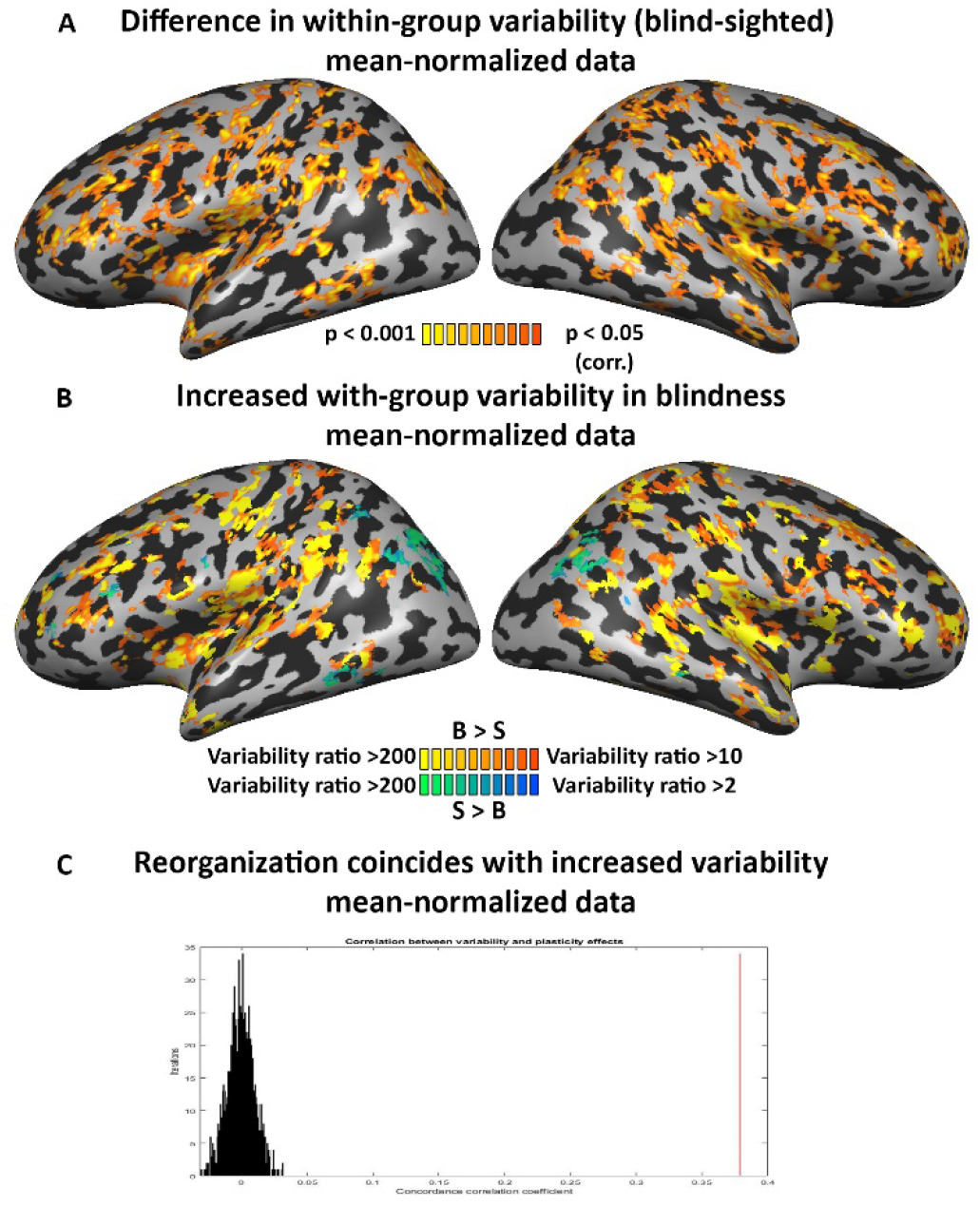
Variability increase in blindness is found regardless of changes to mean RSFC. (**A**) The difference in variability between the groups is significant in various parts of the brain, including in the frontal lobe, even when controlling for the higher mean RSFC values in the blind. (**B**) The blind show increased variability (ratio of blind intra-group variability divided by sighted intra-group variability > 10) in most of the regions differing in their variation between the groups, when controlling for the higher mean RSFC values in the blind. (**C**) Overall across the brain, areas showing changes in RSFC in blindness also show increased variability across blind participants, when controlling for the higher mean RSFC values in the blind. Concordance correlation coefficient was calculated between the RSFC group difference and RSFC change in variability for the V1 seed (red line) and compared to a permutation test (distribution in black).

**Figure S3:**
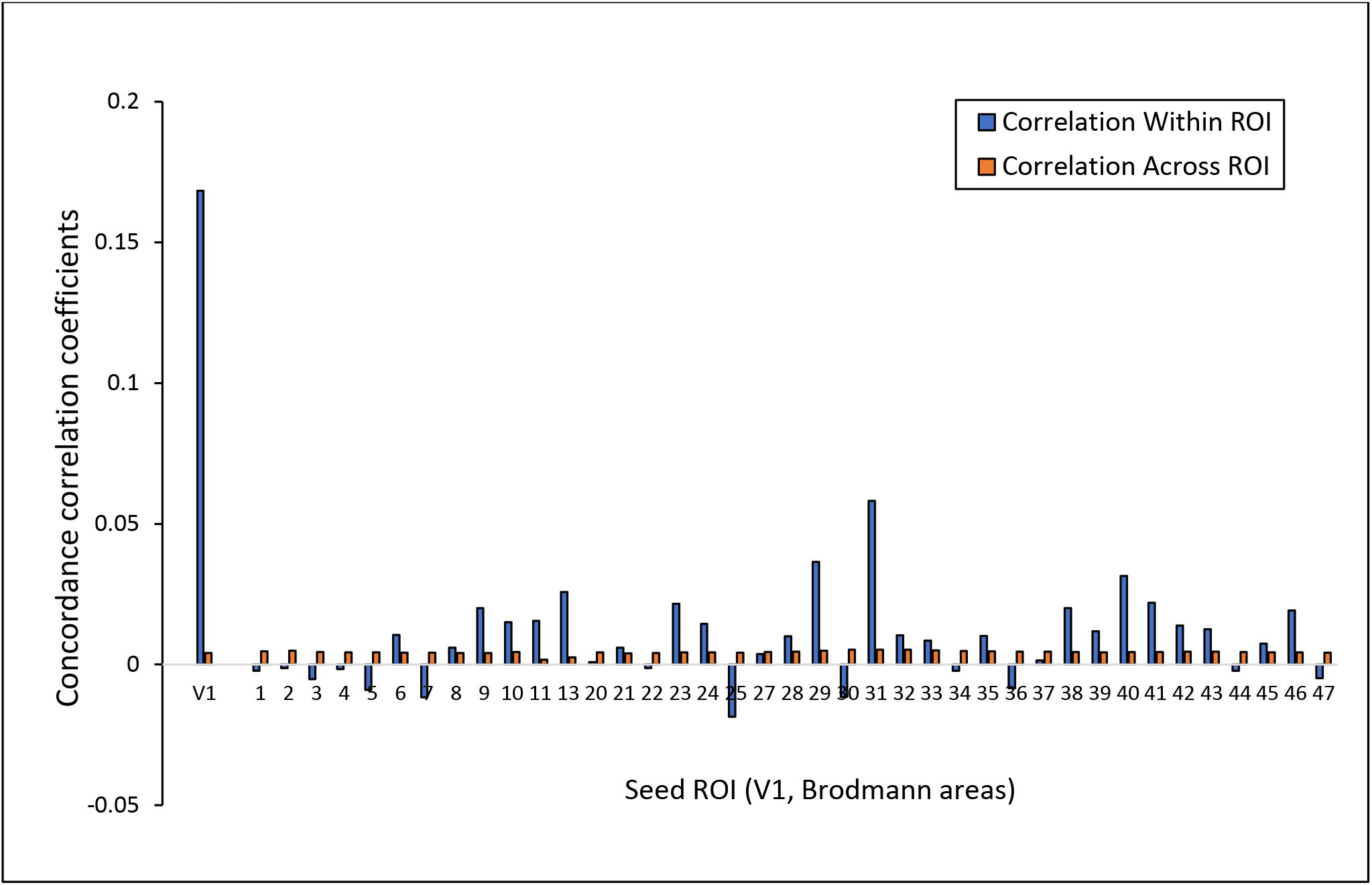
Correlation between variability difference and reorganization is unique to the visual deprived cortex. The correlation (CCC) between reorganization (main effect of group for each seed RSFC; comparable to **Fig. S1A**) and increased variability (Brown-Forsythe test map for each seed RSFC; comparable to **Fig. 1A**) for each control Brodmann area is shown in blue, alongside the value for the V1 seed. For each area, cross-seed correlation (e.g. correlation between the group difference for BA1 and the variability difference of any other Brodmann area; computed in a gray matter mask), was also computed (shown in red). Overall, for all non-visual BAs the within-seed correlation was lower than for V1. Curiously, on average the correlation between each non-visual Brodmann area’s reorganization map and its own variability difference map (blue bars) was slightly higher than the correlated for permuted maps across BAs (red bars), manifesting in a statistical trend (paired t-test, *p*< 0.054; see **Fig. 2D**), and suggesting that more broadly, reorganization manifests in greater variability.

**Figure S4:**
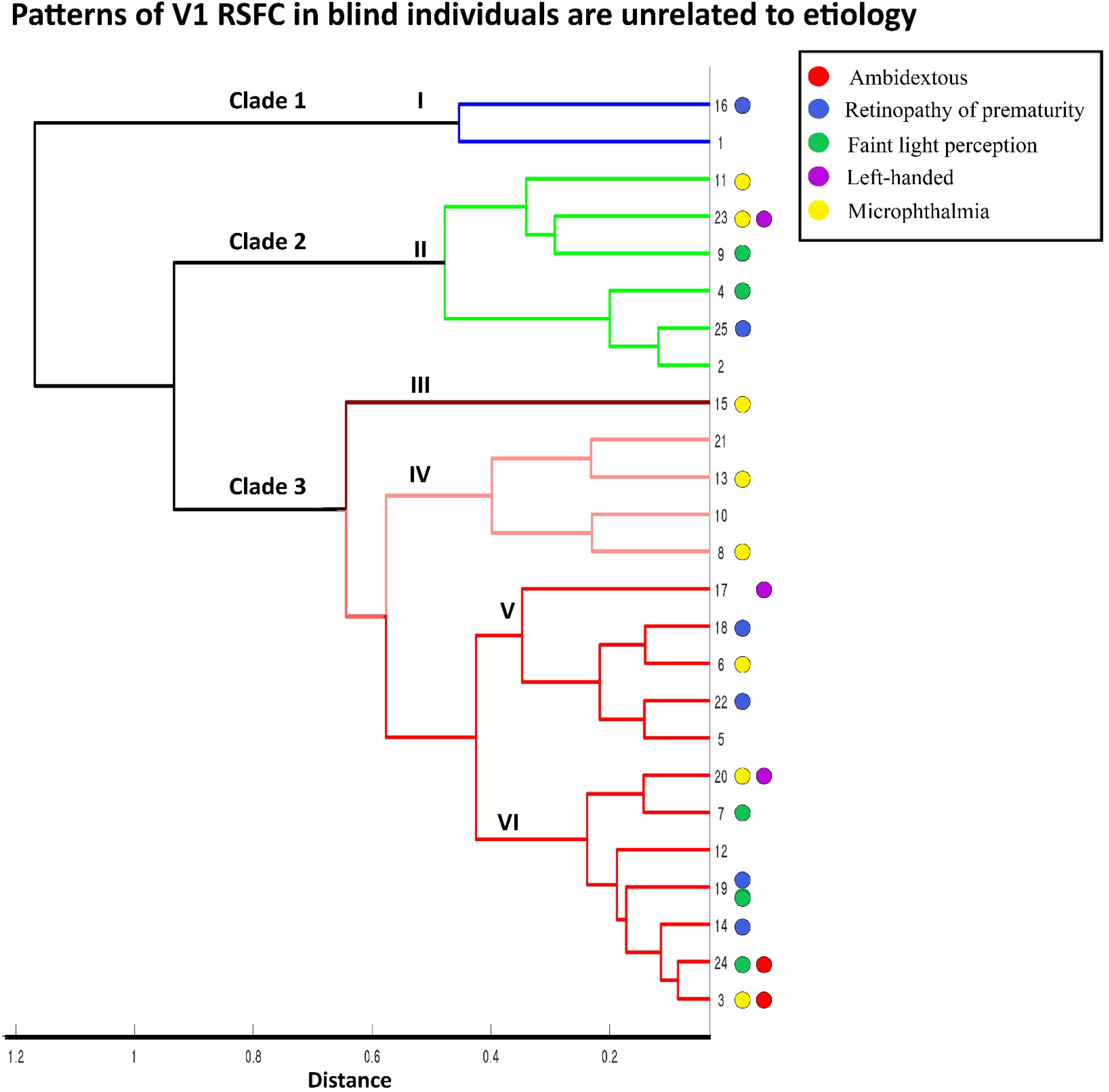
Clustering is not based on blindness etiology. The hierarchical clustering dendrogram depicted in Fig. 3A is repeated, with color indication of frequent blindness etiologies (ROP – blue, microphthalmia – yellow) and unique behavioral traits (ambidextrous individuals – in red, left handedness - in purple and faint light perception – in green). With the exception of the two ambidextrous individuals being clustered together, no other qualitative pattern is evident linking blindness etiology or light perception to the similarity in V1 RSFC profiles. Participants 13 and 20, found on different subclades, are siblings blind due to genetic microphthalmia.

